# ERLIN1 may be involved in DHRS1-induced Change in Lipid Droplet Morphology in HeLa Cell

**DOI:** 10.1101/2023.12.09.570901

**Authors:** Adekunle Toyin Bamigbade, Ololade Omolara Ogunsade, Shimeng Xu, Yaqin Deng, Pingsheng Liu

**Affiliations:** Department of Biological Sciences, Thomas Adewumi University, Oko, Nigeria; Department of Microbiology, Obafemi Awolowo University, Ile-Ife, Nigeria; Institute of Biophysics, Chinese Academy of Sciences, Beijing China

**Keywords:** Short chain dehydrogenase/reductase (SDR), DHRS1, ERLIN1, LD morphology, Lipid droplets

## Abstract

Dehydrogenase/reductase (SDR family) member1, DHRS1, a member of the conserved short chain dehydrogenase/reductase (SDR) superfamily, has been identified in lipid droplets proteome of different cells and tissues. However, until now, little is known about the potential role of DHRS1 on the lipid droplet (LD). Here, we report that DHRS1 localized to the lipid droplet in Huh7 and HeLa cells and ectopic expression of DHRS1 in HeLa cell induced a significant change in the lipid droplet morphology resulting in nearly 2 fold increase both in lipid droplet size and total triacylglycerol level independent of oleic acid treatment. DHRS1 interacted with ERLIN1, a non-caveolae lipid raft-like domain marker, in HeLa cell and ERLIN1 deficient HeLa cells displayed no detectable change in LD morphology. Although ectopic expression of DHRS1-GFP fusion protein in ERLIN1 deficient HeLa cells resulted in fewer GFP-labeled ring structures relative to WT HeLa cell; thus suggesting that ERLIN1 may be involved in regulating DHRS1 protein turnover. Taken together, these data showed that DHRS1 localized to the LD and induced a significant change in LD morphology which may be regulated by ERLIN1.

## INTRODUCTION

Lipid droplet is a dynamic ubiquitous intracellular organelle composed of a core of neutral lipid bounded by phospholipid monolayer with its associated proteins^*1-3*^. Lipid droplets (LDs) provide a store of energy in form of neutral lipids such as triacylglycerol and a reservoir of different classes of lipids which include, but not limited to, cholesteryl and retinol esters essential for signaling and membrane biogenesis. LDs in different tissues also have distinct protein profiles; however, PAT (<u>p</u>erilipin, <u>A</u>DRP and <u>T</u>IP47) family proteins appear conserved in all eukaryote. Members of short chain dehydrogenase/reductase (SDR) superfamily have also been reported in LD proteome^*4*^. SDR is a large family of NAD(P)(H) dependent oxidoreductases; so classified due to the presence of conserved nucleotide-binding Rossmann fold motif characterized by an α/β folding pattern with a central β sheet bound on both sides by 3 α-helices^*5*^. SDR protein family spans all domains of life and its members displayed a number of roles related to intermediary metabolic functions such as xenobiotics, bioconversion of retinoids and lipid metabolisms. For instance, studies from neuroblastoma cell line and *Xenopus laevis*, suggest that DHRS3 negatively regulates retinoic acid level however it is unclear if DHRS3 modulated retinoic acid either by removing its precursor, all-*trans-*retinal (atRAL) or by inhibiting the enzymes converting the latter to all-*trans-*retinoic acid (atRA)^*6,* *7*^. Elsewhere, Hep27 (DHRS2), another member of the SDR family which localizes to the mitochondria, translocates to the nucleus in its mature form and stabilizes p53 by interacting with Mdm2. Thereby suggesting that SDR may modulate tumor suppressor function^*8*^. Cellular distribution of the SDR members is diverse in nature and they sometimes appear to display spatio-temporal localization pattern. Some members of the SDR superfamily have been reported to associate with LD while others are implicated in metabolic diseases. *Saccharomyces cerevisiae* Env9 localizes to the LDs through its C-terminal hydrophobic domain. When overexpressed, Env9 promotes large LD morphology^*9*^. On the other hand, Rdh10 translocates from mitochondria / mitochondrial associated membrane to the LD only when LD biogenesis is stimulated with oleic acid or retinol in COS7 cells^*10*^. Comparative LD proteome from human liver biopsies in normal and non-alcoholic fatty liver disease (NAFLD) condition identified 17β-HSD13 as a novel LD-associated protein up-regulated in NAFLD^*4*^. The LD proteome mass spectrometry also uncovered DHRS1 (dehydrogenase/reductase (SDR family) member 1) as an LD-associated protein^*4*^. DHRS1 was identified from human fetal cDNA library with prominent tissue distribution in liver and kidney by Northern blot analysis^*11*^. However, until now, little is known about the subcellular location/distribution of DHRS1 and its potential role as an LD-associated protein.

In this study, we investigated the potential role of DHRS1 on LD in HeLa cell. Our data reveal that DHRS1 localized to the lipid droplet and ectopic expression of DHRS1 in HeLa cell induced a significant change in the lipid droplet morphology independent of oleic acid treatment. DHRS1 interacted with ERLIN1 in HeLa cell and ERLIN1 deficient HeLa cell displayed no detectable change in LD morphology. Although ectopic expression of DHRS1-GFP fusion protein in ERLIN1 deficient HeLa cells resulted in fewer GFP-labeled ring structures relative to WT HeLa cell; thus suggesting that ERLIN1 may be involved in regulating DHRS1 protein turnover.

## RESULTS

### DHRS1 localizes to the lipid droplets

DHRS1 has been reported in lipid droplet proteome from human liver biopsies samples both in normal and NAFLD patients^*4*^.To further verify subcellular location of DHRS1, DHRS1 was transiently expressed in Huh7 cell lines. Confocal image indicated that DHRS1 formed a ring structure in Huh7 cell line (Fig. 1A). This phenomenon is common to lipid droplet proteins although not exclusive. More so, when FLAG tagged DHRS1 was stably expressed in HeLa cell line, confocal image of FLAG immunofluorescent sample revealed an FITC-labeled ring structure which co-localized with neutral lipid dye, LipidTOX Red (Fig. 1B). From another perspective, HeLa cells were fractionated using OptiPrep to evaluate intracellular distribution of endogenous DHRS1. Cell lysate was adjusted to 30% OptiPrep and layered beneath 20% and 10% OptiPrep in order to allow for soluble cytosolic proteins and buoyant fractions to be separated in a way that buoyant fractions float at the top relative to dense fractions during density gradient centrifugation. According to Western blot results (Fig. 1C), DHRS1 remained enriched in the buoyant fractions 1-3 which co-fractionated with LDs (TIP47, ACSL1 and ATGL) and, in part with ER (fraction 4, Bip and ERLIN1) proteins. Mitochondrial proteins CPT1b, ATP5B and TIM23 were also enriched in fraction 4 as well. Meanwhile DHRS1 was excluded from cytosolic fractions (7-9, GAPDH). These results indicate that DHRS1 associates with LDs and to some extent with ER and mitochondrion and may not be released into the cytosol.

**Figure 1.**
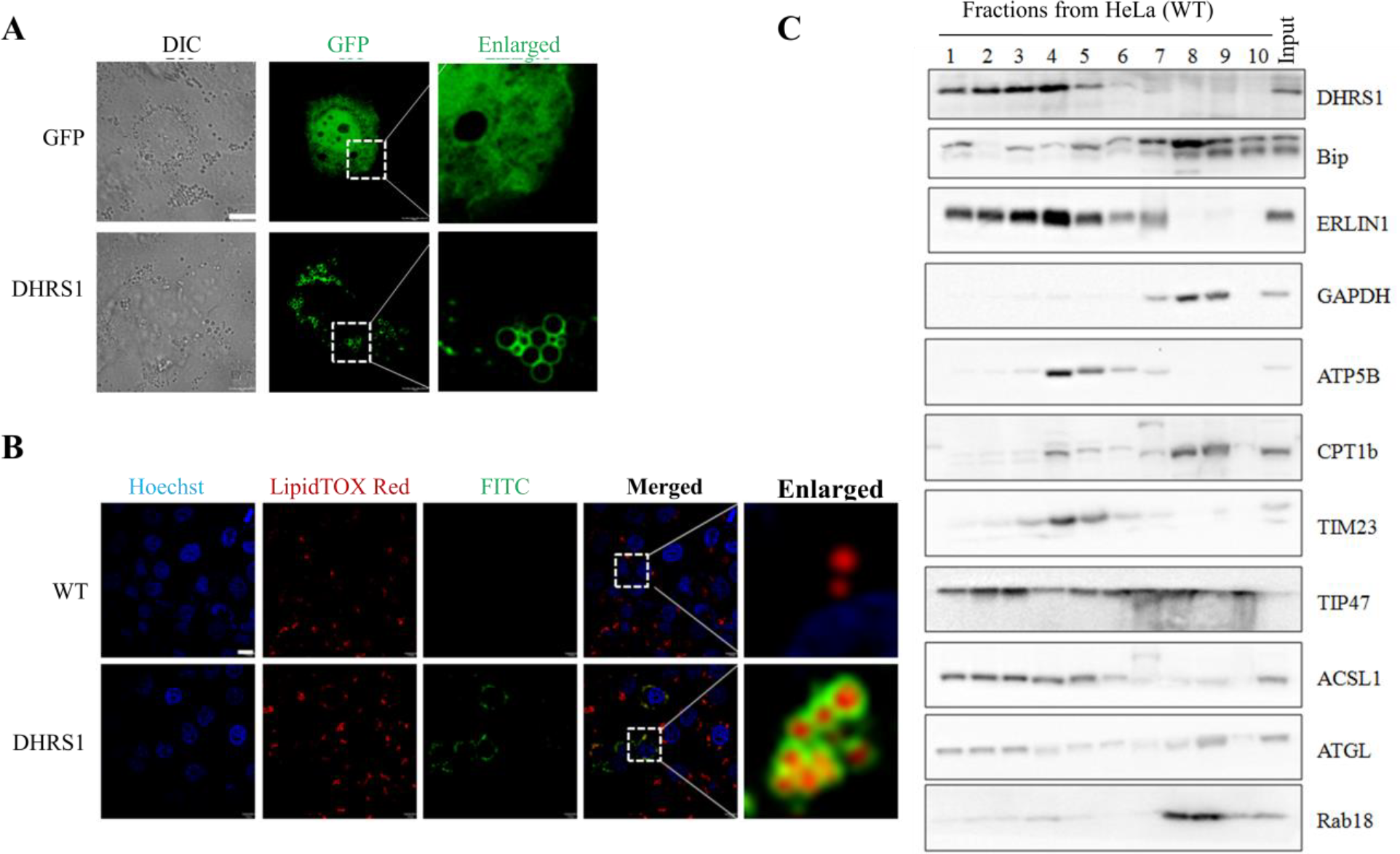
DHRS1 localizes to lipid droplets. (A) GFP tagged DHRS1 was transfected into Huh7 cell line cultured in confocal cell dish with GFP as control. Twenty-four hours (24 h) post-transfection, cells were viewed under confocal microscope and images were captured. (B) HeLa cell line stably expressing DHRS1 tagged with FLAG was cultured on glass cover slips and treated with 100 µM OA supplemented medium for 12 h with WT as control. Cells were fixed with 4% paraformaldehyde for 20 min. Culture medium was removed and cells were washed with PBS. Cells were lysed with 0.1% (v/v) Triton X-100 for 10 min followed by washing with PBS. Lysed cells were blocked with 1% (w/v) BSA and cells were probed with anti-FLAG antibody and FITC conjugated antibody. Cells were washed then incubated with LipidTOX Red and Hoechst (blue) to stain LD and nucleus respectively. Cells were mounted and confocal images were captured. DIC referred to differential interference contrast. Scale bar was set at 10 µm. (C) HeLa cell was cultured to confluence. Medium was removed and cells were washed with PBS and collected in a tube using buffer A supplemented with PMSF. Cells were lysed with N_2_ bomb and 3.0 mg PNS (post nuclear supernatant) was made to 30% OpitPrep. PNS-OptiPrep mixture was loaded into SW40 tube followed by 20% and 10% OptiPrep solutions. Fractionation was carried out at 35,000 rpm for 20 h at 4 °C. Fractions 1 to 10 (that is, top to bottom respectively) were collected, precipitated with trichloroacetic acid and washed with acetone. Proteins were separated on SDS-PAGE and blotted onto PVDF membrane. Following blotting, membranes were probed with indicated antibodies. DHRS1 was enriched in buoyant fractions 1-3 that were also concentrated with TIP47, ACSL1, ATGL representing LDs while fraction 4 enriched in Bip and ERLIN1 (endoplasmic reticulum). ATP5B, CPT1b and TIM23 were enriched in fraction 4 to 5 (mitochondria) while fractions 7 to 9 were rich in GAPDH (cytosol) and Rab18 enriched in fraction 8 to 9.

#### Overexpression of DHRS1 in HeLa resulted in a significant change in LD morphology

To understand potential role of DHRS1 in lipid metabolism, DHRS1 was stably expressed as a FLAG fusion protein in HeLa cells where it induced a significant increase in LD size relative to the control (GFP HeLa cell, Fig. 2A). Hence LD diameter in DHRS1 and GFP cell lines were quantified using ImageJ. In order to quantify LD diameter, DHRS1 and GFP cell lines were treated with and without 100 µM OA for 12 h. Cells were stained with LipidTOX Red and confocal images were captured. In agreement with our observation, LD size increased by nearly 2-fold irrespective of OA-treatment in HeLa cell stably expressing DHRS1 relative to GFP stable cell line (Fig. 2B). Therefore, we hypothesized that increased LD size might have resulted from triacylglycerol (TAG) accumulation. To verify this, total TAG level in DHRS1 stable cell line was evaluated relative to GFP with protein level as a denominator in both cell lines. Interestingly, TAG level in DHRS1 stable cell line increased by 2-fold independent of OA-treatment (Fig. 2C). Furthermore, thin layer chromatography (TLC) results of HeLa cell stably expressing DHRS1 and GFP control treated with and without OA as highlighted above also indicated a 2-fold increase in TAG level in DHRS1 cell lines independent of OA treatment (Fig. 2D). Taken together, these data suggests that DHRS1-induced change in LD morphology may have resulted from increased cellular TAG level.

**Figure 2.**
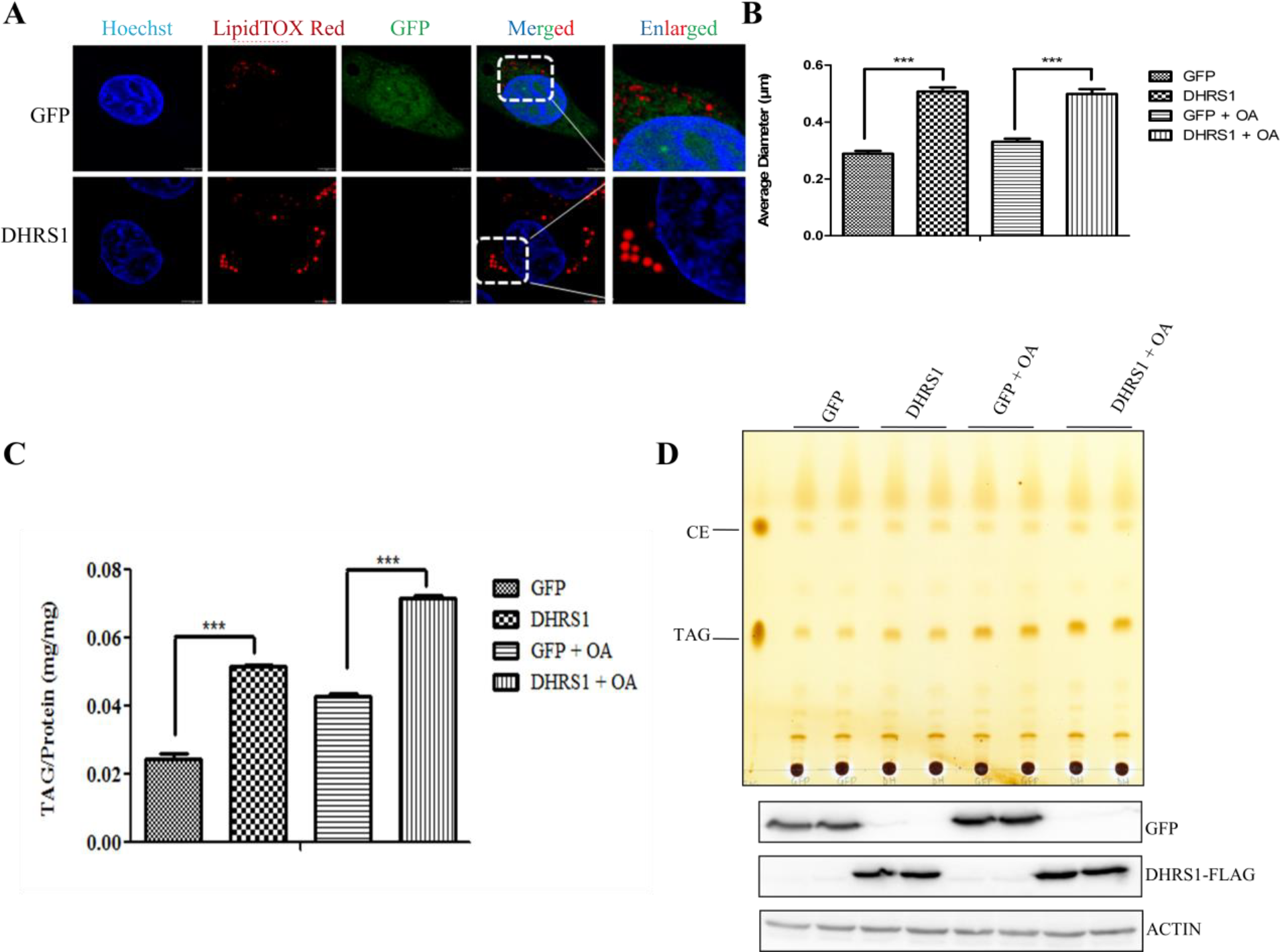
Overexpression of DHRS1 induces large LD phenotype in HeLa cell, in part, due to increase in cellular TAG level. (A) HeLa cells stably expressing DHRS1 protein were cultured on glass cover slips and treated with 100 µM OA for 12 h where GFP served as control. Cells were fixed with 4% (v/v) paraformaldehyde then washed with PBS. Cells were treated with LipidTOX Red and Hoechst (blue) to stain LDs and nucleus respectively. (B) Average of LD diameter from twenty cells (20) each from DHRS1 and GFP cell lines with or without OA treatment for 12 h were counted using ImageJ on a scale bar of 2 µm. *** p< 0. 0001. (C) Total triacylglycerol increased significantly by nearly 2-fold in DHRS1 cell line relative to GFP stable cell line independent of OA treatment. Indicated stable cell lines (DHRS1 and GFP control) were cultured to confluence in 6-well dishes and treated with and without 100 µM OA for 12 h. Cells were collected in phosphate buffered saline pH 7.4 containing 1% (v/v) Triton X-100 and sonicated. Aliquot of cell lysate was subjected to TAG (triacylglycerol) analysis with protein as internal control. All TAG experiments were in 4 replicates with*** p< 0. 0001. (D) Neutral lipid analysis by TLC. Lipids from respective cell lines with indicated treatment were separated by TLC with a solvent of n-hexane-diethyl ether-acetic acid (80:20:1, v/v/v). TLC plate was then stained in iodine vapor after being dried at room temperature. Where CE is cholesterol ester; TAG, triacylglycerol. Western blots of GFP (negative control) and DHRS1-FLAG indicating stable expression of respective proteins and ACTIN served as protein loading control.

### ERLIN1 interacted with DHRS1 in HeLa cell

Protein-protein interaction studies have lent itself as an essential tool to understand a gene function. To unravel a possible mechanism by which DHRS1 induces change in LD morphology and TAG accumulation, DHRS1 cell line was subjected to FLAG immunoprecipitation (see Methods). Precipitate was eluted with sample buffer and resolved in SDS-PAGE followed by silver staining. Detected distinct band (Fig. 3) was submitted to Q Exactive MS. Table 1 indicated the list of proteins identified from our MS data. ALDOA, ERLIN1 and GNAI3 were selected for further study due to their high unique peptide number. These three genes were fused with Myc/His at their C-termini and transfected into DHRS1 stable cell line for 24 h followed by FLAG immunoprecipitation. At first, Western blot of DHRS1 precipitate displayed no detectable interaction between DHRS1 and any of the three proteins-ALDOA, ERLIN1 and GNAI3) (not shown). Of course, this might be due to a weak interaction between DHRS1 and interacted protein(s) drawing from the barely detectable level of the identified band (Fig. 3). Hence, DHRS1 cell line was again transfected with ALDOA, ERLIN1 and GNAI3 followed by treatment with 1 mM DSP, a bifunctional crosslinker, and FLAG iimmunoprecipitation. Of the three selected genes, only ERLIN1 co-interacted with DHRS1 (Fig. 4). Interestingly, WT HeLa cell fractionation (Fig. 1 C) had earlier indicated that DHRS1 co-fractionate with ERLIN1 (indicated by both LD and ER markers). Although protein co-fractionation does not necessarily mean the two proteins interact. Both IP and MS data thus suggest that DHRS1 interacts with ERLIN1.

**Table 1.** List of DHRS1 interacting proteins identified by mass spectrometry.

| Band Number | Protein Name | Peptide Number | MW (kDa) | Accession |
| --- | --- | --- | --- | --- |
| 1 | DHRS1 | 15 | 33.9 | Q96LJ7 |
|  | TIM50 | 6 | 39.6 | Q3ZCQ8 |
|  | GNAI3 | 6 | 40.5 | P08754 |
|  | ALDOA | 6 | 39.4 | P04075 |
|  | ERLIN1 | 6 | 38.9 | O75477 |
|  | MP2K3 | 4 | 39.3 | P46734 |
|  | FPPS | 2 | 40.5 | P14324-2 |
|  | AIMP2 | 2 | 35.3 | Q13155 |

**Figure 3.**
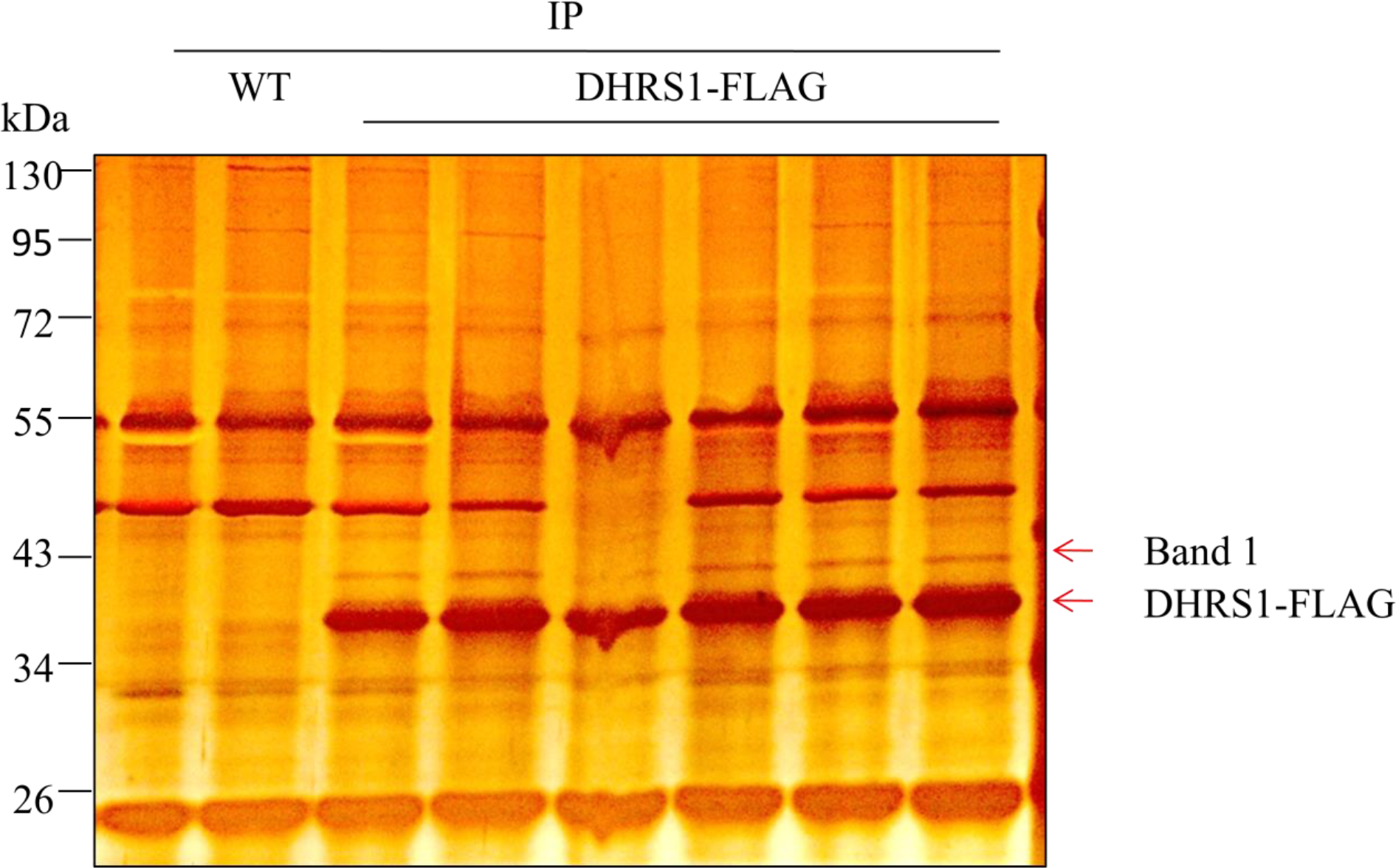
Identification of unique DHRS1 precipitated proteins in HeLa cell. HeLa cell line expressing FLAG tagged DHRS1 was subjected to immunoprecipitation with GFP as control. Precipitated proteins were eluted in sample buffer. Proteins were separated on SDS-PAGE and stained with silver nitrate. Indicated band 1 was cut, in-gel digested with trypsin and submitted for mass spectrometry.

**Figure 4.**
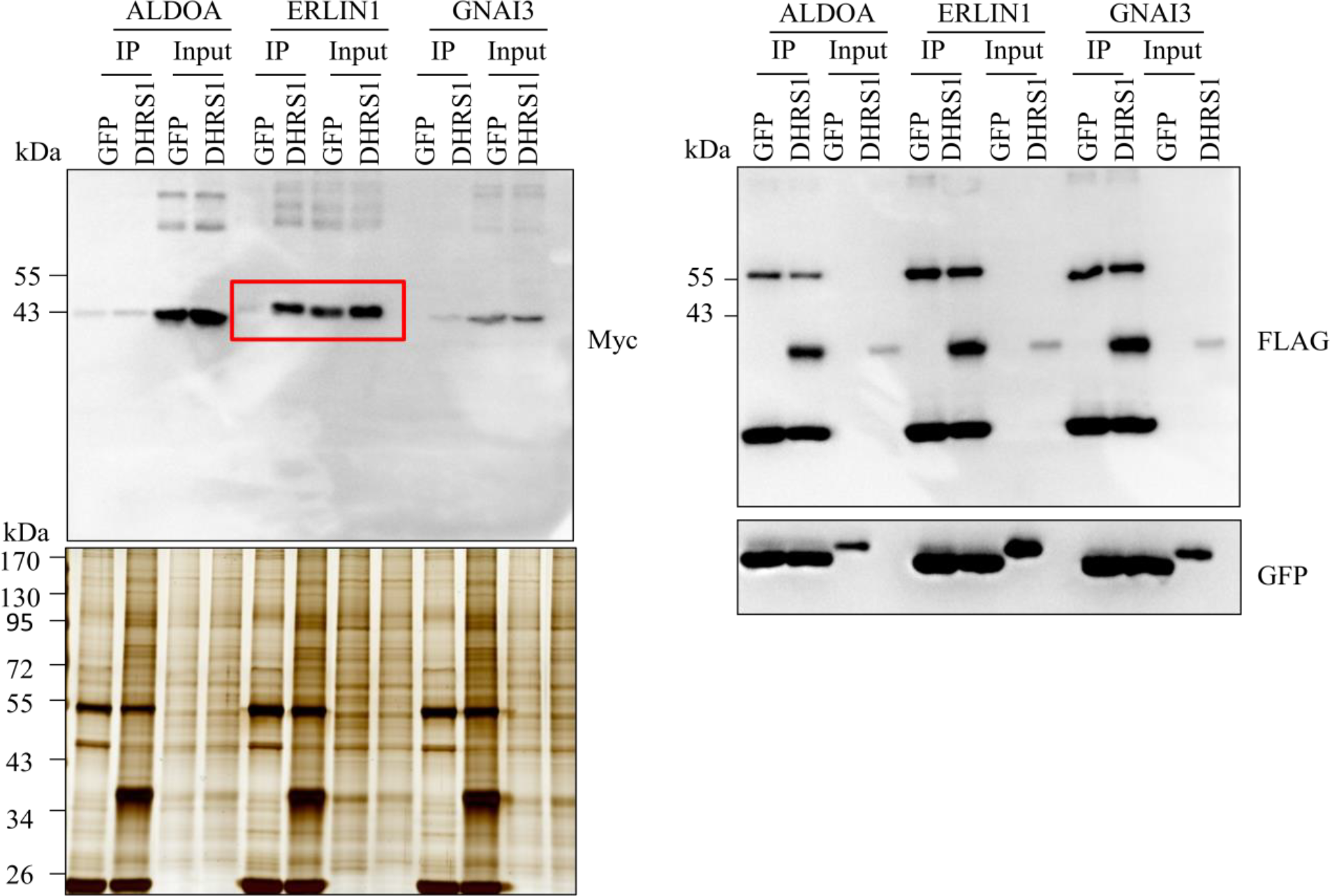
DHRS1 interacted with ERLIN1 in HeLa cell. Three of the identified proteins (Table 1) were cloned into pCDNA3.1 plasmid and transfected into DHRS1 stable cell line as ALDOA, ERLIN1 and GNAI3 fused with Myc tag. FLAG immunoprecipitation was carried out 24 h post-transfection and precipitated proteins were eluted in sample buffer. Protein sample was separated on SDS-PAGE and blotted onto PVDF membrane and probed with Myc- and FLAG-antibodies. Prior to FLAG immunoprecipitation, cells were treated with 1 mM DSP cross linker for 30 min and reaction was quenched by Tricine buffer. GFP and silver stained gel served as control. IP, immunoprecipitation.

#### No detectable change in LD morphology in ERLIN1 deficient HeLa cells

Since ERLIN1 interacted with DHRS1 as identified by mass spectrometry and validated through co-IP in DHRS1 stable cell line, ERLIN1 may be a key player in DHRS1-induced change in LD morphology earlier observed. Thus, ERLIN1 was knocked out in WT HeLa cell by CRISPR-Cas9 endonuclease technique. ERLIN1 KO was verified by PCR analyses of genomic DNA from two representative clones (TIC6 and T2C19) which were derived from disruptions of ERLIN1 ORF at exon1 and exon2; and by Western blot analysis with WT as negative control (Fig. 5). Then, WT (HeLa) and ERLIN1 KO clones (T1C6 and T2C19) were prepared for confocal image after treatment with 100 µM OA for 12 h. Confocal image results (Fig. 6) indicated no detectable change in LD morphology between WT and ERLIN1 deficient HeLa cells. This data suggest that ERLIN1 KO has no detectable effect on LD morphology in HeLa cell.

**Figure 5.**
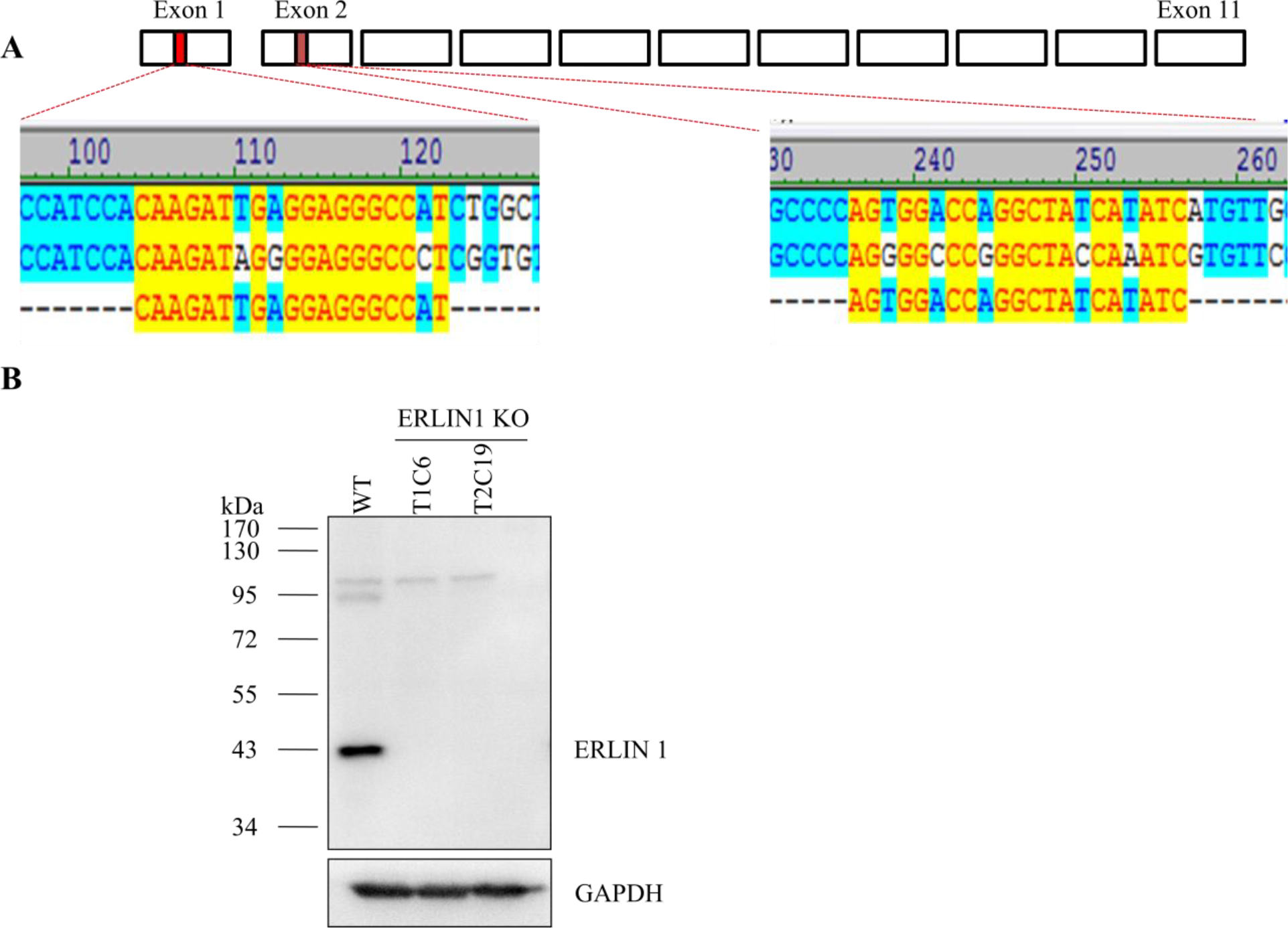
ERLIN1 knockout by CRISPR-Cas9 technique in HeLa cell. (A) HeLa cells were transfected with sgRNA (single guide RNA) targeting Cas9 endonuclease encoded in pX260a plasmid to indicated locations on exon1 and exon2 in the genomic locus encoding for ERLIN1. Regions highlighted in yellow represents sequence results of disrupted ERLIN1 gene. (B) ERLIN1 knockout in HeLa by sgRNA1 (target1, T1) and sgRNA2 (target2, T2) was verified by Western blot where GAPDH served as loading control and WT served as negative control. T1C6 indicated target1 clone 6 while T2C19 indicated target2 clone 19; and KO indicated knockout.

**Figure 6.**
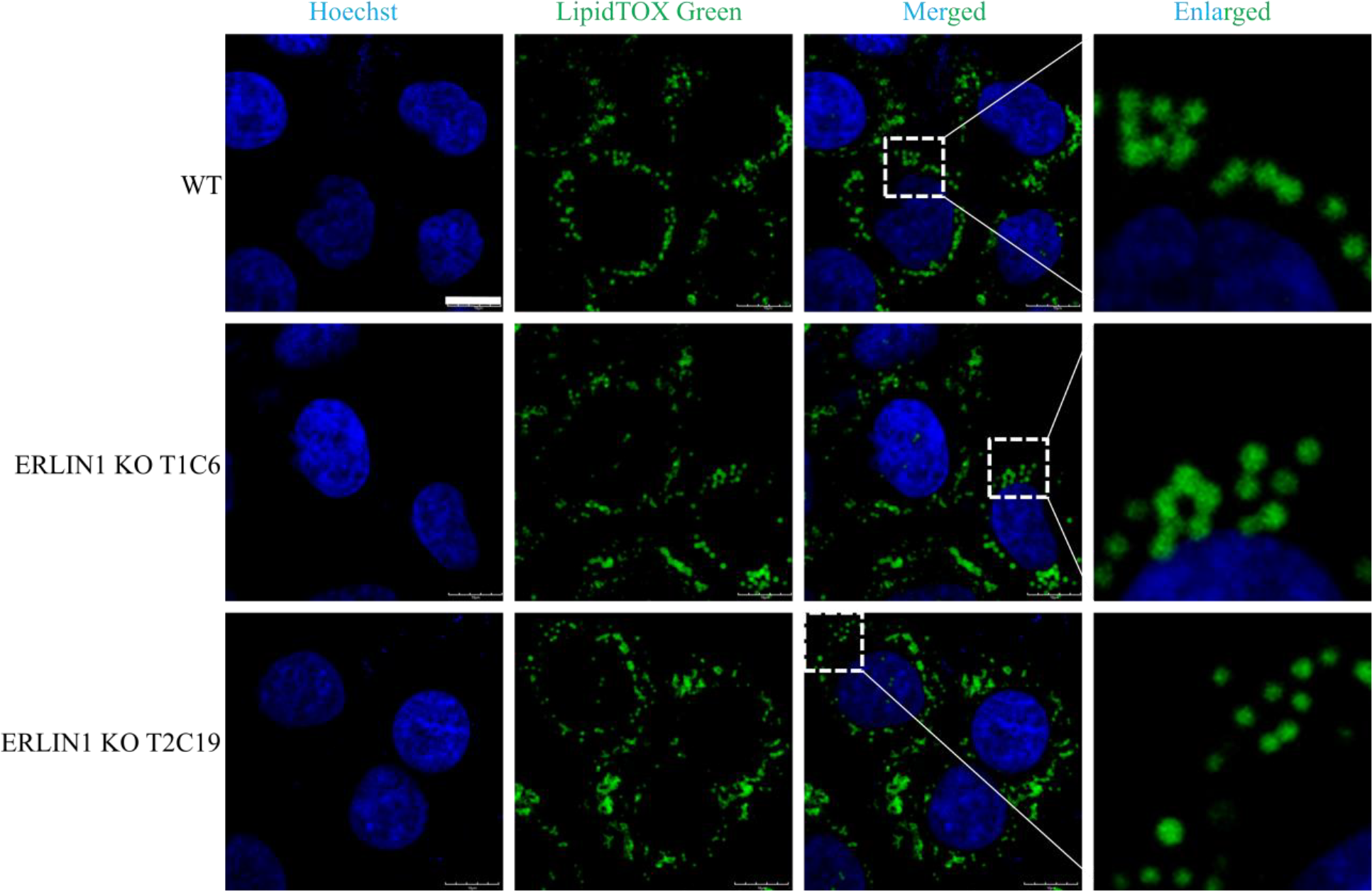
No detectable change in lipid droplet morphology in ERLIN1 deficient HeLa cells. HeLa cells harboring ERLIN1 knockout and WT HeLa cells were separately cultured on glass cover slips and treated with 100 µM OA for 12. After this treatment, cells were fixed and incubated with Hoechst (blue) and LipidTOX Green to stain nucleus and lipid droplets respectively. Confocal images were captured and scale bar was set at 10 µm.

#### ERLIN1 deficient HeLa cells stably expressing DHRS1 displayed fewer LDs

Next, DHRS1 was stably expressed as a GFP fusion protein in HeLa (WT) and HeLa cells harboring ERLIN1 KO (T1C6 and T2C19) and confocal images of cells in respective groups were captured. Interestingly, confocal image (Fig. 7) indicated that ERLIN1 deficient HeLa cells contained fewer sparsely distributed GFP-labeled ring structures compared to WT.

**Figure 7.**
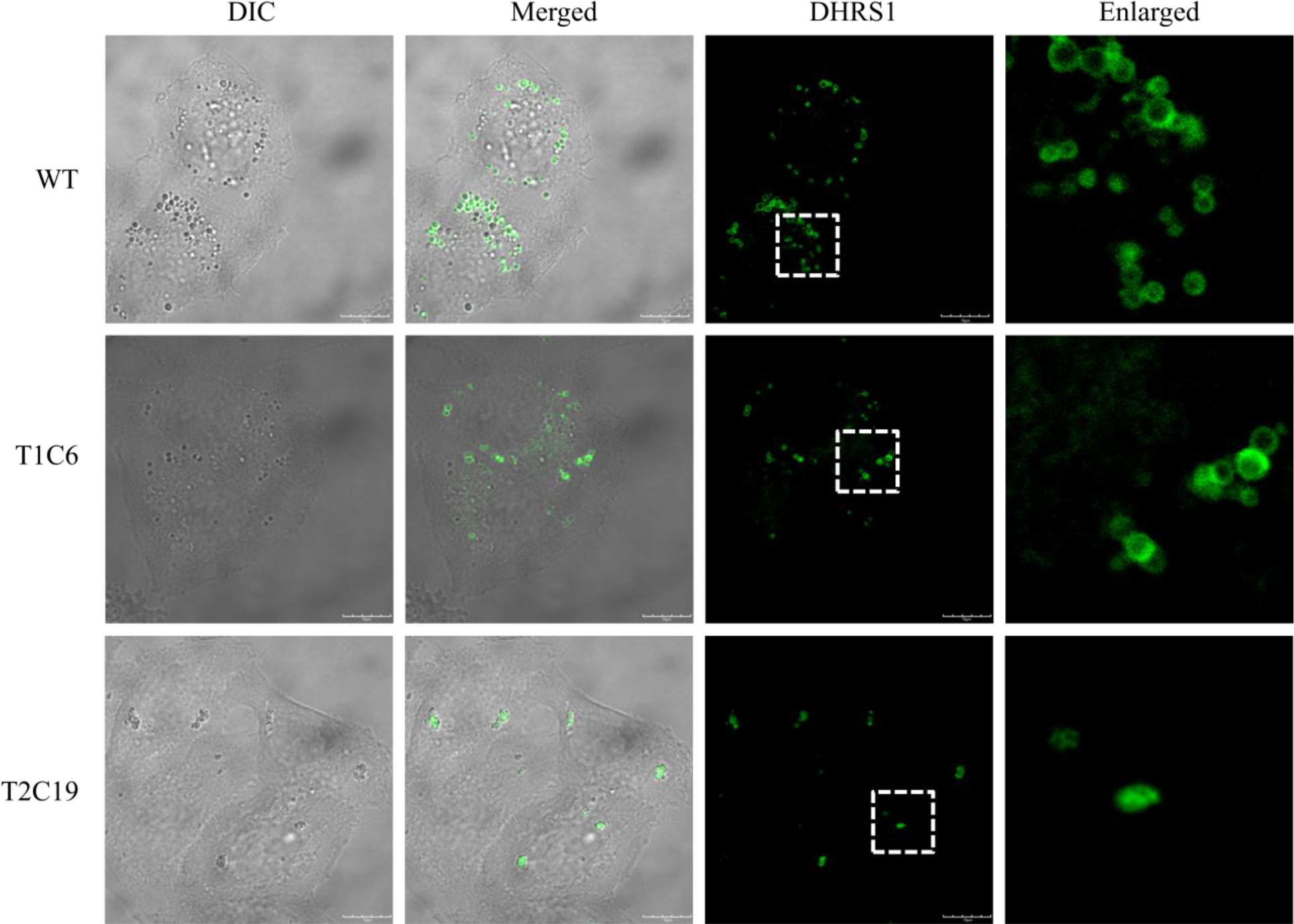
ERLIN1 knockout results in fewer GFP-labeled ring structures in DHRS1 stable cell. DHRS1-GFP was stably expressed in WT (HeLa) and HeLa cells harboring ERLIN1 KO (T1C6 and T2C19). Indicated stable cells were cultured on glass cover slips and confocal images were captured. WT indicated wild type; T1C6 indicated target1 clone 6; T2C19 indicated target2 clone 19. Scale bar was set at 10 µm.

Tentative data suggest that ERLIN1 may be involved in stabilizing DHRS1 protein although further experiments are required to validate this finding.

## DISCUSSION

Members of the SDR superfamily reportedly distributes to different organelles within the eukaryotic cells. *Saccharomyces cerevisiae* Env9 localized to the LD while Rdh10 both localized to the LD or mitochondria/mitochondrial associated membrane depending on cellular status^*9, 10*^. On another hand, DHRS4 location is isoform dependent, isoform 1 localized to the peroxisome while isoform 7 localized to the nucleus^*12, 13*^. In this study, we showed that DHRS1 localized to the LDs in HeLa, and Huh7 cells by confocal microscope mages of GFP fusion protein and immunofluorescence of FLAG tagged DHRS1. More so, DHRS1 majorly co-fractionated with LD and, in part, with ER and mitochondria marker proteins. Furthermore, series of repeated LD isolation followed by Western blot analyses showed that DHRS1 localized to the LD in HeLa cell. A recent report, however, indicated that DHRS1 localized to the ER independent of the position of the epitope tag used; interestingly, the work was also carried out in HeLa cell^*14*^. First, DHRS1 appears to display multiple subcellular localization (LD and ER) based on the cell fractionation **result**. Although, confocal microscope image may not be sufficient to conclude the subcellular location of a protein; immunostaining and immunoblot analyses using monoclonal antibody could be more specific for antigen binding and subcellular localization evaluation. Also, pieces of information gathered from the Uniprot database, suggest that post-translational modification would remove the initiator methionine from human DHRS1 thus making the N-terminal region of the protein unsuitable for epitope fusion analyses. However, cell status and other yet to be identified factors may be involved in subcellular protein distribution and or processing. Representative members of the lipid raft proteins such as flotillin-1 and stomatin have been identified as LD proteins in CHO K2 and MDCK cells respectively ^*15, 16*^. More so, flotillin-1 association with LD is OA-dependent similar to caveolins ^*17, 18*^. Intriguingly, ERLIN1, another member of the lipid raft proteins, co-fractionated with DHRS1 in HeLa cell line thus suggesting that ERLIN1 may be an LD-associated protein.

Studies on *Saccharomyces cerevisiae* Env9 protein, an ortholog of human retinol dehydrogenase 12 (RDH12), have shown that ectopic expression of Env9 positively modulated LD size, we hypothesized that DHRS1 may play a similar role in LD dynamics. In agreement with our prediction, HeLa cell with a stable expression of DHRS1 displayed a significant increase in LD size. This is further supported by a significant increase in cellular triacylglycerol level which is independent of exogenous fatty acid supplement. Large LD phenotype and TAG accumulation appear to be a recurring theme in SDR superfamily members implicated in lipid metabolism^*9*^ for instance, DHS-3 in *C*.*elegans* and its human homolog type11 hydroxysteroid (17-β) dehydrogenase (HSD17β11) in HeLa^*19*^.

SDR proteins which target to the LD display large LD phenotypes when overexpressed. In this study, we adopted protein-protein interaction approach to probe the mechanism underpinning this phenomenon. Rounds of immunoprecipitation of FLAG-tagged DHRS1 coupled with mass spectrometry followed by co-IP experiments of myc-tagged ALDOA, ERLIN1 and GNAI3 implicated ERLIN1 as a DHRS1-interacting protein in HeLa cell. More so, stable expression of DHRS1-GFP in ERLIN1 deficient HeLa cells indicated fewer GFP-labeled ring structures relative to control. ERLIN1/KEO4p/SPFH1 is a marker of the non-caveolar lipid raft-like domain intrinsic to the ER^*20*^. ERLIN1 also forms a functional complex with ERLIN2 in the ER. It binds cholesterol and is involved in the regulation of SREBP1. ERLIN1 has been associated with metabolic diseases such as non-alcoholic fatty liver disease and haemorrhagic stroke^*21*^. Only recently, ERLIN1 was reported to display an increased copy number variation (CNV) in a patient with coronary artery disease. While ERLIN1 is closely associated with lipid rafts, other reports indicated that ERLIN1 is essential for ER-associated degradation (ERAD)-a mechanism that selectively degrades misfolded proteins^*22, 23*^.To the best of our knowledge, until now, no report has neither suggests interaction of ERLIN1 protein with DHRS1 nor its co-fractionation with lipid droplet proteins. Although, the role of ERLIN1 interaction with DHRS1 in our study is not completely understood, ERLIN1 may, however, serve to facilitate neutral lipid transfer to the LD or regulate DHRS1 protein turnover. Interestingly, overexpression of DHRS1-GFP in ERLIN1 deficient HeLa cells suggests that ERLIN1 may serve to stabilize DHRS1 protein.

In conclusion, DHRS1 localized to the LD and ectopic expression of DHRS1 in HeLa cell resulted in significant increase in LD size and cellular TAG level and ERLIN1 may be important in regulating DHRS1 protein turnover. Although further study is required to understand the potential mechanism by which ERLIN1 regulate DHRS1 protein and also verify if ERLIN1 plays any significant role in DHRS1-induced TAG accumulation and LD size increase in HeLa cell.

## Author Contributions

P. L. supervised the project. A. T. B., O. O. B. S. X., and Y.D. conducted the experiments. P. L., A. T. B., designed the experiments and analyzed the data. A. T. B. wrote the manuscript.

## MATERIALS AND METHODS

Mouse anti-GFP was purchased from Santa Cruz and rabbit anti-Myc from Cell Signaling Technology. Mouse anti-FLAG M2 was purchased from Sigma. For other antibodies (see Table S1). The Colloidal Blue staining kit, Hochest 33342 and LipidTOX Red were from Invitrogen. Pierce BCA Protein Assay kit from Thermo Scientific and TAG kit from Biosino Bio-technology and Science Inc. pCDNA3.1 was from Invitrogen while pEGFP-N1 and pQCXIP were obtained from Clontech.

### Cell Culture

HeLa cell lines (American Type Culture Collections, Manassas, VA) were maintained in DMEM (Macgene Biotech., Beijing) supplemented with 10% FBS (Gibco), 100 U/ml penicillin and 100 mg/ml streptomycin at 37 °C, 5% CO_2_.

### Construction of Plasmids

Coding sequences used in this study were amplified from HeLa cell line cDNA library and cloned into respective vectors. All epitope tags-GFP, Flag, and Myc-were fused to the C-terminal part of the respective proteins. Primer pairs are listed in Table S2. In order to establish a stable DHRS1-FLAG / GFP cell line, corresponding cDNAs were cloned into pQCXIP plasmid and HeLa cells were infected with retrovirus expressing these proteins followed by puromycin screening until monoclones were established and verified by immunofluorescence microscopy. In order to generate ERLIN1 deficient cells, single guide RNA (sgRNA) was ligated into pX260a vector encoding CRISPR-Cas9 endonuclease and transfected into WT HeLa cell. Single clones of HeLa cells deficient in ERLIN1 were screened by Western blot and insertion/deletion mutation at indicated ERLIN1 genomic locus was validated through PCR analysis.

### Confocal Microscopy

HeLa cells stably expressing DHRS1 protein were cultured on glass cover slips and treated with 100 µM OA for 12 h where GFP served as control. Cells were fixed with 4% (v/v) paraformaldehyde (PFA) for 20 min rinsed with phosphate buffer saline (PBS) followed by 1:1,000 (v/v) LipidTOX Red in PBS washed and subsequently treated with 1:1,000 (v/v) Hoechst in PBS for 10 min before being mounted with cover slips. For immunofluorescent analysis, after fixing with 4% PFA, cells were lysed with 0.1% (v/v) Triton X-100, blocked with 1% (w/v) BSA and probed with mouse anti-FLAG antibody and subsequently, goat anti-Mouse IgG conjugated with FITC. LipidTOX Red and Hoechst treatment were the same as stated above.

### Immunoprecipitation

HeLa cell line stably expressing DHRS1-Flag and GFP were cultured to confluence in 100 mm dishes. The cells were rinsed with cold PBS, collected into DNase-free Eppendorf tubes in 1 mlTETN solution containing 150 mM NaCl supplemented with 100x protease inhibitor. The lysate was incubated for 30 min and centrifuged at 4 °C for 10 min at 17,000*g*. Clarified supernatant was incubated with anti-FLAG conjugated sepharose beads and eluted with 2x sample buffer containing β-mercapto-ethanol after washing with TETN buffer containing 500 mM, 250 mM and 150 mM NaCl. The sample was subjected to silver and colloidal blue staining. DHRS1 precipitated protein band was subjected to Q Exactive mass spectrometry for identification.

### Lipid Droplet Isolation

LDs were purified using the method described previously with modifications^*24*^. Briefly, HeLacell treated with 100 µM oleic acid for 12 h were scraped and collected after 3 rinses with ice-cold PBS. Then all the cells were transferred to 50 mL buffer A (25 mM tricine pH 7.6, 250 mM sucrose) containing 0.5 mM PMSF. After centrifugation at 3,000 *g* for 10 min the cell pellets were re-suspended in 10 ml buffer A containing 0.5 M PMSF and were incubated on ice for 20 min. The cells were then lysed by N_2_ bomb (750 psi for 15 min on ice). The cell lysate was centrifuged at 3,000 *g*, the post-nuclear supernatant (PNS) fraction was collected and loaded into a SW40 tube and was overlain with buffer B (20 mM HEPES, pH 7.4, 100 mM KCl, and 2 mM MgCl_2_). The sample was centrifuged at 250,000*g* for 1 h at 4 °C. The white band containing LDs at the top of gradient was collected into a 1.5 ml centrifuge tube. The sample was centrifuged at 20,000*g* for 10min at 4°C and then the underlying solution was carefully removed. The droplets were gently re-suspended in 200 μl buffer B. This procedure was repeated four times. The lipid extraction and the protein precipitation were carried out by chloroform/acetone (1:1, v/v) treatment followed by centrifuging the sample at 20,000*g* for 30 min at 4 °C. The protein pellet was then dissolved in 2× sample buffer (125 mM Tris Base, 20% glycerol, 4% SDS, 4% β-mercaptoethanol and 0.04% bromophenol blue).

### In-gel Digestion of Proteins

The gel bands containing the protein sample were manually excised. Each of the protein bands was then digested individually as below. The protein bands were cut into small plugs, washed twice in 200 μl of distilled water for 10 min each time. The gel bands were dehydrated in 100% acetonitrile for 10 min and dried in a Speedvac for approximately 15 min. Reduction (10 mM DTT in 25 mM NH_4_HCO_3_ for 45 min at 56 °C) and alkylation (40 mM iodoacetamide in 25 mM NH_4_HCO_3_ for 45 min at room temperature in the dark) were performed, followed by washing of the gel plugs with 50% acetonitrile in 25 mM ammonium bicarbonate twice. The gel plugs were then dried using a speedvac and digested with sequence-grade modified trypsin (40 ng for each band) in 25 mM NH_4_HCO_3_ overnight at 37 °C. The enzymatic reaction was stopped by adding formic acid to a 1% final concentration. The solution was then transferred to a sample vial for LC-MS/MS analysis

### LC-MS/MS Analysis

All nano LC-MS/MS experiments were performed on a Q Exactive (Thermo Scientific) equipped with an Easyn-LC 1000 HPLC system (Thermo Scientific). The labeled peptides were loaded onto a 100 μm id×2 cm fused silica trap column packed in-house with reversed phase silica (Reprosil-Pur C18AQ, 5 μm, Dr. Maisch GmbH) and then separated on an a 75 μm id×20 cm C18 column packed with reversed phase silica (Reprosil-Pur C18AQ, 3μm, Dr. Maisch GmbH). The peptides bounded on the column were eluted with a 75-min linear gradient. The solvent A consisted of 0.1% formic acid in water solution and the solvent B consisted of 0.1% formic acid in acetonitrile solution. The segmented gradient was 4–12% B, 5 min; 12–22% B, 50 min; 22– 32% B, 12 min; 32-90% B, 1 min; 90% B, 7min at a flow rate of 300 nl/min.

The MS analysis was performed with Q Exactive mass spectrometer (Thermo Scientific). With the data-dependent acquisition mode, the MS data were acquired at a high resolution 70,000 (m/z 200) across the mass range of 300–1,600 m/z. The target value was 3.00E+06 with a maximum injection time of 60 ms. The top 20 precursor ions were selected from each MS full scan with isolationwidth of 2 m/zforfragmentationin the HCD collision cell withnormalized collision energy of 27%. Subsequently, MS/MS spectra were acquired at resolution 17,500 at m/z 200. The target valuewas 5.00E+04with a maximum injection time of 80 ms. The dynamic exclusion time was 40s. For nano electrospray ion source setting, the spray voltage was 2.0 kV; no sheath gas flow; the heated capillary temperature was 320°C. For each analysis, 2 µg peptides were injected and each sample was measured in duplicate.

### Protein Identification

The raw data from Q Exactive were analyzed with Proteome Discovery version1.4 using Sequest HT search engine for protein identification and Percolator for FDR (false discovery rate) analysis against a Uniprot Human protein database. Some important searching parameters were set as following: trypsin was selected as enzyme and two missed cleavages were allowed for searching; the mass tolerance of precursor was set as 10 ppm and the product ions tolerance was 0.02 Da; the methionine oxidation was selected as variable modifications; the cysteinecarbamidomethylation was selected as a fixed modification. FDR analysis was performed with Percolator and FDR <1% was set for protein identification. The peptides confidence was set as high for peptides filter.

### Western Blot

Proteins were separated on SDS-PAGE gel and blotted onto PVDF membrane. The membrane was blocked in 5% defatted milk and probed with respective antibodies. Protein bands were detected by incubating membrane in enhanced chemiluminescent substrate after incubating with indicated second antibodies.

### Cell Fractionation

HeLa cells were cultured until the cell density was 90%. Cells were collected and rinsed with cold PBS. Cells were collected in solution A supplemented with PMSF. Cell sample was centrifuged at 3,000*g* for 10 min. Supernatant was discarded and cell pellet re-suspended with solution A. Cell sample was lysed under pressure and centrifuged at 1,000*g* for 10 min. Post-nuclear supernatant (PNS) was analyzed for protein concentration using the BCA method. PNS-containing 30% OptiPrep was loaded in SW40 tube followed by 20 % OptiPrep and 10% OptiPrep. The sample was centrifuged at 35,000 rpm for 20 h at 4 °C. Fractions were collected from top to bottom and treated and protein precipitated with TCA and cold centrifuged at 15,000 rpm for 10 min. The supernatant was discarded. Subsequently acetone was added and the precipitate centrifuged as above. Meanwhile after each acetone washing, pellet was re-suspended by sonication. Samples were air-dried for 10 min and dissolved in 2x sample buffer and denatured at 95 °C for 5 min. Samples were centrifuged to clarify protein and used for Western blot analysis.

#### Immunofluorescence

Cells were cultured to around 70% cell density on glass slides, fixed with paraformaldehyde in DMEM for 20 min. and rinsed with cold PBS. The cells were permeabilized with Triton X-100 in PBS for 10 min. Cells were incubated with BSA for 1 h (blocking). The cells were then treated with mouse monoclonal anti-FLAG antibody with and left to incubate for 1 h. Following incubation with primary antibody, cells were rinsed with PBS. Goat anti-mouse antibody conjugated to FITC was applied to the cells for another 1 h. Subsequent steps in sequential order include, washing of cells with PBS, treatment of cells with LipidTOX Red for 30 min and Hoechst for 10 min. The slides were mounted by immersion oil and confocal images were captured.

### Triacylglycerol Quantitation

Indicated cell lines were sub-cultured into 6-well dishes grown to confluent and treated with or without 100 µM OA for 12 h. Cells were rinsed with cold PBS, aspirated and collected with 1% (v/v) Triton X-100 in PBS. Cellular triacylglycerol level was subsequently analyzed with TAG kit and denominated with total protein evaluated with BCA kit according to manufacturer’s instruction.

### Thin Layer Chromatography Analysis

Indicated cell lines were sub-cultured into 6-well dishes grown to confluence and treated without or with 100 µM OA for 12 h. Cells were rinsed with cold PBS, aspirated and collected with 1% (v/v) Triton X-100 in PBS. TAG was extracted from cell samples using chloroform and methanol in ratio 2:1. Resulting supernatant was dried under a stream of nitrogen and analyzed with TLC, developing solvent of hexane-diethyl ether-acetic acid (80:20:1, v/v/v). TLC plates were stained in iodine vapor after drying at room temperature.

### Statistical Analyses

Data were presented as mean ± SEM unless specifically indicated. The statistical analyses were performed using GraphPad Prism 6. Comparisons of significance between groups were performed using unpaired Student t-tests by Quickcalc.

